# Resident and Engrafting *E. coli* Populations Expand Through Dissimilar Pathways in the Inflamed Gut

**DOI:** 10.64898/2026.04.29.721686

**Authors:** Manuela Roggiani, Jun Zhu, Mark Goulian

## Abstract

Intestinal inflammation increases the abundance of *Enterobacteriaceae* in the gastrointestinal tract by several orders of magnitude. These population expansions, or blooms, are associated with disease progression and have been suggested to exacerbate intestinal pathologies in some settings. Murine studies have shown that during the early stages of *Escherichia coli* colonization, i.e., during engraftment, inflammation enhances fitness through processes that depend on Moco, an enzyme cofactor found in a variety of oxidoreductases that consists of molybdenum coordinated by a pterin molecule. Using a murine commensal *E. coli* isolate and a DSS-induced colitis model in mice, we investigated whether Moco is also important for blooms of *E. coli* that are part of the resident microbiota, that is, for *E. coli* that have engrafted well before the onset of inflammation. We show that resident wild-type and Moco^-^ *E. coli* exhibit comparable expansions in response to inflammation, indicating that, in this context, Moco-dependent processes such as nitrate respiration or formate oxidation were not important for inflammation-induced blooms. We find that Moco is important, however, for *E. coli* colonization in the absence of inflammation, suggesting that alternative respiratory pathways or other Moco-dependent processes are necessary for *E. coli* colonization of a healthy murine gut. Our findings demonstrate that the mechanisms underlying inflammation-induced blooms can depend on the temporal relationship between engraftment and inflammation, and also highlight the importance of considering colonization stage in identifying and interpreting the factors that affect the fitness of microbes colonizing the intestine.

## Importance

Inflammation in the intestine disrupts the bacterial populations that inhabit this environment. One characteristic feature of this disruption is the large increase in abundance—blooms—of *Escherichia coli* and related species. These blooms have received considerable attention because they have been associated with disease progression of various forms of colitis. Previous work in mice identified respiratory pathways important for the fitness of *E. coli* that are introduced into the intestine around the same time that inflammation is induced. However, there is a common assumption that the same pathways account for blooms of *E. coli* that have colonized well before the onset of inflammation. Here, we show an example in which this assumption is incorrect. Therefore, one may need to distinguish between early and late colonization stages to understand some of the mechanisms underlying inflammation-induced blooms. This distinction is also likely to be important for many other aspects of microbial colonization of the intestine.

Fitness determinants of microbes in the intestine are likely to depend on colonization stage. Upon invading the GI tract, successful colonizers must transit myriad environments, with varying nutrients, stressors, and other properties, before successfully establishing a foothold in a suitable niche. The temporal organization of this process is poorly described, but one can at least distinguish between the earliest stages, where the colonizers are still in the process of engrafting, and much later stages, where they have reached a stable or quasi-stable state and can be considered members of the resident microbiota. Many of the physiologic requirements for growth, survival, and retention in the intestine are likely to be different for these early and late stages, and therefore, many of the genes most important for fitness are likely to differ as well. This view is supported by longitudinal studies, including competitions, transposon sequencing, and transcriptomic experiments (1, 2). For the same reasons, a microbe’s response to intestinal perturbations, such as a change in diet or the onset of inflammation, could also depend on colonization stage. In this work, we show that this is indeed the case for the role of molybdenum cofactor (Moco) in the response of an *E. coli* isolate to dextran sodium sulfate (DSS)-induced inflammation in the murine intestine.

Intestinal inflammation has a profound effect on the microbiome and can cause the abundance of *E. coli* and other *Enterobacteriaceae* to increase by several orders of magnitude (3-5). These population expansions, or “blooms,” have been associated with disease progression and might also play a causal role in some intestinal pathologies (6-9). Although much of the microbial physiology and ecology underlying inflammation-induced blooms has not been fully characterized, studies focused on *E. coli* have identified Moco to be a key fitness determinant for the colonization of the inflamed murine intestine (10). *E. coli* contains nineteen Moco-dependent enzymes, which function in various metabolic pathways such as electron transport chains and repair reactions (11). Through analysis of specific mutants, two such processes—nitrate respiration and formate oxidation—have been shown to be important for *E. coli* colonization of DSS-treated mice as well as in other inflammation models (10, 12, 13). Furthermore, these studies showed that inflammation increases intestinal nitrate and formate levels. Most mouse studies comparing the fitness of wild-type *E. coli* to that of mutants defective in Moco synthesis or in a Moco-dependent pathway have focused on intestinal inflammation occurring early in the colonization process (i.e., inflammation was induced either prior to, concurrently with, or shortly after *E. coli* inoculation.) Here, we explore whether Moco-dependent processes necessarily play a similar role in blooms of *E. coli* that are part of the resident microbiota prior to the onset of inflammation.

For this study, we used a murine commensal *E. coli* isolate, MP1 (14), and constructed a Moco**-** derivative by deleting *moaA*, which encodes the enzyme catalyzing the first committed step of Moco biosynthesis (11). Since Moco is required for nitrate respiration and *E. coli* cannot ferment glycerol (11, 15), we verified that the Δ*moaA* strain was unable to grow anaerobically on glycerol as a carbon source in minimal medium supplemented with nitrate (Fig. S1A). We then repeated the experiments that established Moco is a fitness determinant for *E. coli* colonization of the inflamed intestine (10). As in this previous study, wild-type and Δ*moaA* strains were co-inoculated into mice either four days after (Figs. 1A; S2A) or 24 hours before (Fig. 1B; S2B) DSS was introduced into the drinking water to induce acute colitis. In these competitions, the wild-type strain showed a significant advantage over Δ*moaA*, consistent with previous observations of Moco’s importance for *E. coli* fitness when colonization was initiated in temporal proximity to the induction of colonic inflammation with DSS (10).

**Figure 1.**
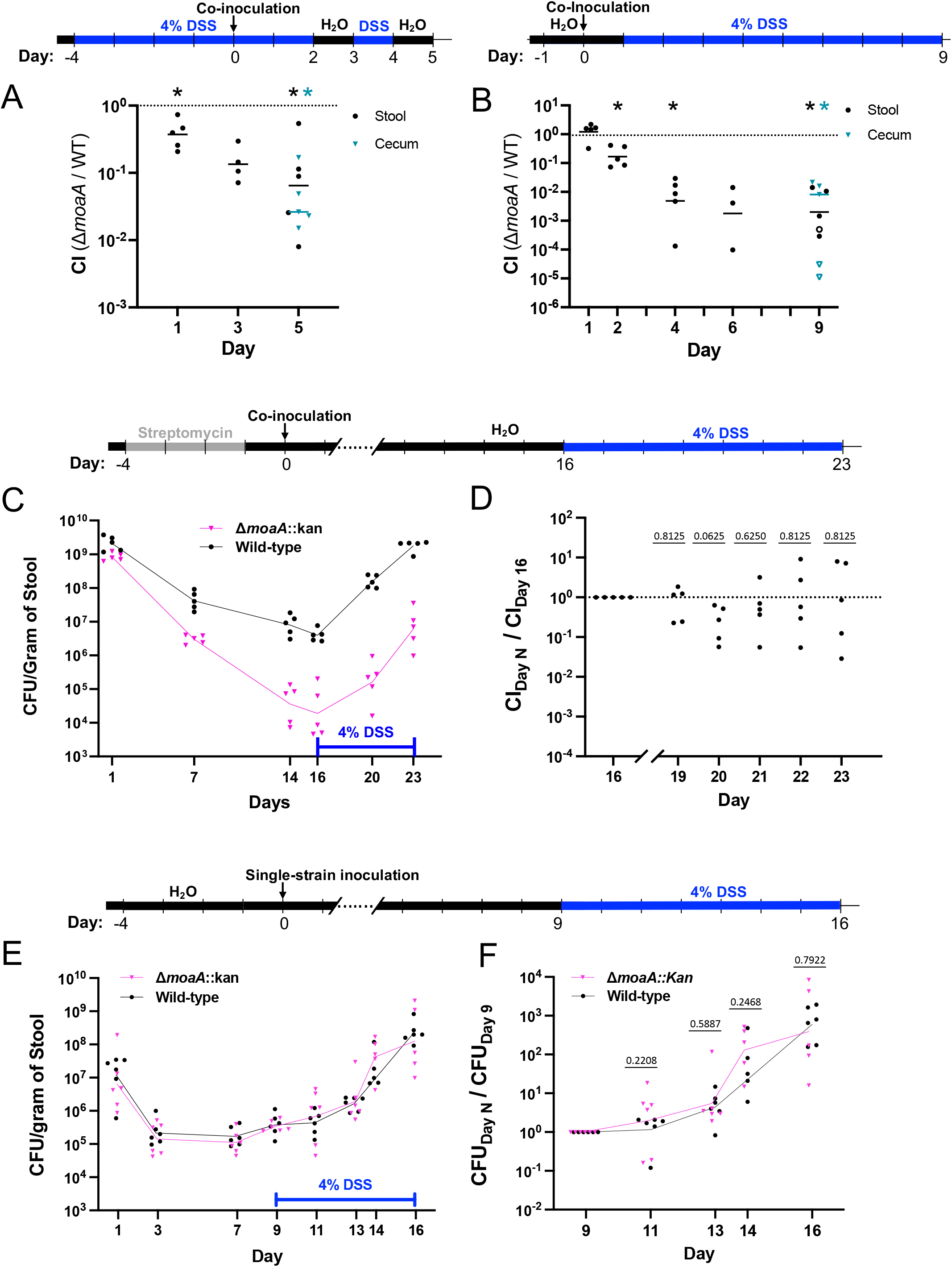
Fitness effects of DSS on Δ*moaA E. coli* depend on the timing of DSS treatment relative to inoculation. (A,B) Competitions between Δ*moaA* and wild type in which mice were treated with DSS prior to (A) or 24 hours after (B) inoculation, as indicated and as described previously (10). Competitive index (CI) is equal to: Δ*moaA* CFU/wild-type CFU at the indicated time point divided by the corresponding ratio for the inoculum, where CFU denotes colony-forming units. The open circles indicate values for which the Δ*moaA* counts were set equal to the detection limit because no Δ*moaA* colonies were detected on plates. Asterisks denote p<0.05 based on the Wilcoxon matched-pairs signed-rank test – one-tailed to test Δ*moaA <* wild type, which was observed in (10). The corresponding CFU for these experiments are shown in Fig. S2. (C) Competitions between Δ*moaA* and wild type in which mice were inoculated 24 hours after streptomycin pretreatment and given DSS for seven days, beginning on day 16 following inoculation. Points denote the stool CFU from individual mice, and lines connect the geometric means. (D) The relative fold change in abundance following DSS treatment for the data shown in (C). The relative fold change on day N of DSS treatment is defined to be the ratio of the competitive index, CI, on day N to the CI just prior to DSS treatment on day 16: CI_Day N_ ⁄CI_Day 16_. Note that this ratio satisfies 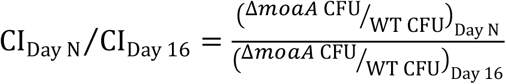. The numbers above the data points denote p-values from a Wilcoxon matched-pairs signed-rank test comparing wild type and Δ*moaA*.(E) Single-strain colonizations of either Δ*moaA* or wild type. In these experiments, mice were not pre-treated with antibiotics, and were given DSS for seven days beginning on Day nine. (F) Ratio of the stool CFU on the indicated day relative to the stool CFU just prior to DSS treatment on Day 9 for the data shown in (E). The numbers above the data points denote p-values from a Wilcoxon signed-rank test. For each of the competition experiments in (A-C), five mice were used and were co-housed in the same cage. For the single-strain colonization experiments (E), a total of 6 mice were used for each *E. coli* strain and housed in 3 cages (2 mice per cage). All DSS-treated mice showed evidence of weight loss and disease activity indices of at least 3 by the end of the experiments. In addition, histological analysis of colonic sections indicated clear signs of inflammation (Fig. S4).

Next, we tested whether Moco plays a similar role in *E. coli* blooms when inflammation is induced well after *E. coli* have engrafted in the intestine. Mice were pre-treated with streptomycin and then co-colonized with Δ*moaA* and wild-type strains. We waited for stable colonization (two weeks following inoculation) before introducing DSS into the drinking water. In two repeats of this experiment performed on different dates, *E. coli* counts declined more slowly (Fig. S3A,B) and stayed well above the colonization levels that we typically observe for stable colonization of mice by *E. coli* MP1 (Fig. 2), (14). This difference between (Fig. 1C) and the two repeats in (Fig. S3A,B) likely reflects differences in the recovery of the microbiota following streptomycin pre-treatment. Therefore, for the experiments in (Fig. S2A,B), the mice were gavaged with a slurry of stool from untreated mice three weeks after inoculation to hasten the decline in CFU to levels we observe in long-term colonization (Fig. 2). In all three experiments, following DSS administration, fecal counts of wild-type *E. coli* increased and, by the seventh day, reached levels that were higher by a factor of 10^2^-10^3^ over those measured just prior to DSS treatment (Fig. 1C; S3A,B). Notably, the levels of the Δ*moaA* strain also increased in response to DSS. Although there was considerable variability among mice in the relative increases of wild-type versus Δ*moaA* counts, the data indicate a similar propensity for expansion of both strains, as assessed by the competitive index of Δ*moaA* versus wild type at time points during DSS treatment, normalized by the competitive index immediately before DSS treatment was started (Fig. 1D; S3A,B).

**Figure 2.**
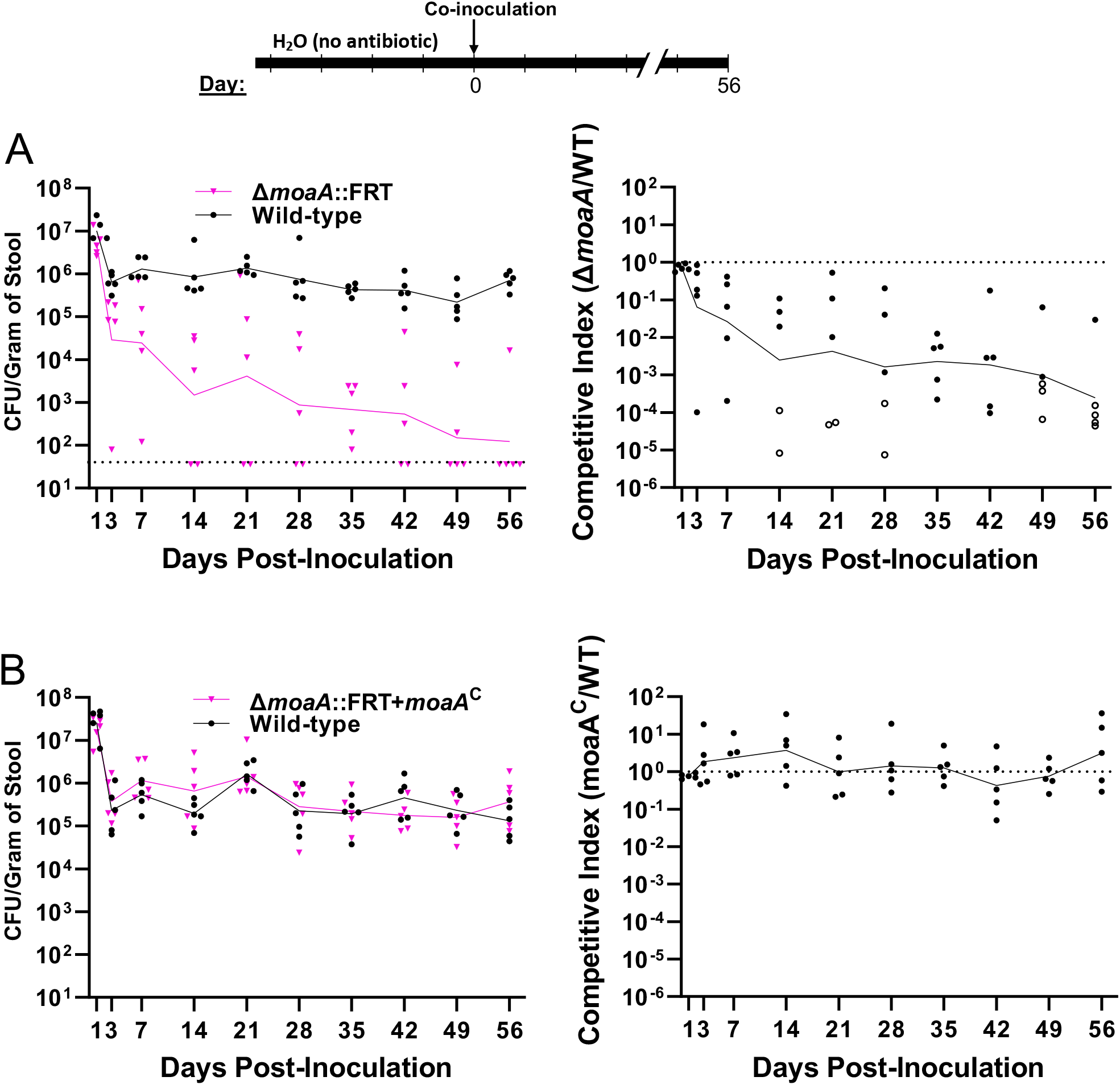
Competitions between wild-type and Δ*moaA E. coli* in mice without antibiotic pre-treatment (A) and complementation of Δ*moaA* (B). For these experiments, we used a Δ*moaA* strain in which the kanamycin resistance gene was removed, leaving an in-frame deletion in *moaA*, in order to minimize the chance of a polar effect on downstream genes in the *moaABCDE* operon and to facilitate complementation. The dotted line (A, left) denotes the detection limit, and the open circles (A, right) indicate competitive indices for which the Δ*moaA* counts were set equal to the detection limit because no Δ*moaA* colonies were detected on plates. The notation *moaA*^C^ indicates that the *moaA* gene was inserted at the *att*_Tn7_ site. Competitive index is (Δ*moaA* CFU)/(wild-type CFU) at the indicated time point divided by the corresponding ratio for the inoculum. In (B) competitive index is defined similarly, except with *moaA*^*c*^ CFU instead of Δ*moaA* CFU. For each experiment (A and B), the five mice were housed in a single cage.

We also tested whether Δ*moaA* and wild type showed similar DSS-induced blooms when mice were engrafted with only a single strain. For these experiments, we took advantage of the fact that *E. coli* MP1 stably colonizes SPF C57BL/6 mice without antibiotic pre-treatment. Mice were colonized with either wild type or Δ*moaA* without streptomycin pretreatment, and colonization levels stabilized to ~10^5^-10^6^ cfu per gram of stool within 72 hours. After nine days, DSS was added to the drinking water, and we again observed that Δ*moaA* and wild type showed comparable blooms (Figs. 1E,F).

In the above competition experiments, levels of the Δ*moaA* strain declined markedly relative to wild type during the several weeks prior to DSS treatment (Figs. 1C; S3A,B). Since it has been reported that streptomycin treatment of mice can produce a mild inflammatory response that promotes *E. coli* growth through nitrate respiration (16), we tested whether this decline is a result of intestinal perturbation by streptomycin. In competition experiments without streptomycin pretreatment (and without DSS treatment), Δ*moaA* still showed a colonization defect relative to wild type (Figure 2A). In addition, insertion of *moaA* at an ectopic site in the chromosome complemented this deletion both in mouse competitions (Fig. 2B) and *in vitro* (Fig. S1B). These results indicate that the absence of Moco causes a competitive fitness defect in the healthy mouse intestine.

Taken together, our findings indicate that the mechanisms underlying *E. coli* MP1 population expansions in response to DSS-induced colitis can differ for early and late stages of colonization. When inflammation was present during the engraftment process, i.e., before or very shortly after *E. coli* was introduced into the GI tract, we found that Moco was a key fitness determinant (Figs. 1A,B), consistent with previous reports (10). On the other hand, for *E. coli* that had already engrafted and were effectively part of the resident microbiota when inflammation commenced, Moco’s absence had no detectable effect on the population expansion. Thus, at least for the *E. coli* strain and colonization model used here, nitrate respiration and formate oxidation, both Moco-dependent processes, are not important for blooms of *E. coli* that have colonized well before DSS treatment, even though they are important when *E. coli* is engrafting in an inflamed intestine (10, 12). We note, however, that a prior study showed that tungstate, which inhibits Moco synthesis by competing with molybdate, suppresses the DSS-induced bloom of resident *Enterobacteriaceae* in mice. This suggests that Moco-dependent processes are important for the expansion of at least some resident *Enterobacteriaceae*, including possibly some *E. coli* strains. We also note that the results for our colonization model do not rule out a role for respiration since some terminal reductases, such as those that reduce oxygen or fumarate, do not use Moco. Indeed, aerobic respiration has been suggested to play a role in *E. coli* population expansions in response to colitis, at least for early colonization stages (12, 17).

Although we found that deleting *moaA* did not impact DSS-induced blooms of resident *E. coli*, it did lower *E. coli* colonization fitness in competitions with wild type when inflammation was not induced (Figs. 1C; S3A,B; 2A). This finding may indicate a role for Moco-containing respiratory enzymes in the colonization of a healthy intestine. However, it could also reflect the importance of other Moco-dependent pathways in supporting colonization, such as pathways associated with detoxification, repair, or nutrient acquisition (18-22).

Our results also highlight the importance of considering colonization stage when assessing the effects of genetic and environmental perturbations on bacterial fitness in the gut. The factors that influence the success of an initial colonization, i.e., engraftment, and those that influence persistent colonization are both important for the microbial ecology of the intestine. However, colonization dynamics are poorly understood, even for relatively simple systems. More work exploring the temporal organization of colonization is thus needed to identify the mechanisms that control the composition and host interactions of intestinal microbiota.

## Acknowledgements

We would like to thank Aaron Hecht, Josie Ni, and Gary Wu for discussions and comments on the manuscript and Nathan Fanzone for histopathology analysis. This work was supported by NIH grants R21AI125814 (MG and JZ), R01AI157106 (JZ), R35GM139541 (MG).

