## Supplementary Information for "Resident and Engrafting *E. coli* Populations Expand Through Dissimilar Pathways in the Inflamed Gut"

**Materials and Methods**
**Table S1. List of Strains**
**Table S2. List of Primers**
**Figs. S1-S4**

### Materials and Methods

*Bacterial Growth*

Bacterial strains were cultured at 37 °C in Miller lysogeny broth (LB) (Fisher Scientific), or M9 minimal medium (23) supplemented with 2 mM $FeSO_4$ and 0.2% glucose or 0.4% glycerol. When indicated, $KNO_3$ was added to anaerobic cultures in M9 to a concentration of 40 mM. For the preparation of electrocompetent cells, bacteria were cultured in SOB (Amresco, Solon, OH) with the appropriate antibiotic when necessary. Antibiotics for selection of recombinants or for the maintenance of plasmids were added to bacterial cultures and to LB agar plates at the following concentrations: Kanamycin 50 µg/ml, Ampicillin 50 µg/ml. To induce fluorescent protein expression on agar plates for determining colony-forming units, tetracycline was added to LB agar to a final concentration of 15 µg/ml

*Strain Construction*

The *moaA* deletion (Δ*moaA*) was constructed by recombineering as described previously (14). The *E. coli* strain MP1 carrying the helper plasmid pKD46 was electroporated with a PCR product made from the Δ*moaA*::[FRT-*kan*-FRT] KEIO collection strain JW0764, and primers *moaA*-KEIO-F and *moaA*-lower. The primers were designed to exploit regions of sequence identity flanking *moaA* in MP1 and JW0764, which comprised the 131 bp upstream of the *moaA* start codon, and 124 bp downstream of the stop codon. The *moaA* deletion/insertion was moved via $P1_{vir}$ transduction into strain MP13 by selection for kanamycin resistance. To ensure that the locus $att_{\lambda}$::[ *tetR* Φ(*tetA*-*gfp*)] in MP13 was not eliminated by the transduction (*moaA* is near $att_{\lambda}$), the kanamycin resistant colonies obtained by transduction were screened on tetracycline plates and gfp positive colonies were identified.

The *moaA* complementation strain was constructed by chromosomal integration of a single copy of *moaA* at the Tn7 attachment site of strain MP302. This strain was derived from MP241 by removal of the Kanamycin cassette with pCP20 (24). The moaA gene, including roughly 500 bp of sequence upstream of the start codon was isolated by PCR with the primers *moaA*-XmaI-U1 and *moaA*-EcoRI-L1 and cloned into the *Xma*I and *EcoR*I restriction sites of pUC18R6KT-mini-Tn7T-Km (25), between the neomycin cassette and Tn7-L arm. The resulting plasmid pMR138 and helper plasmid pTNS2 (25) were coelectroporated into strain MP302. The neomycin-resistant clones were checked for insertion at the correct site, and the inserts were sequenced.

*Animal Studies*

Animal studies were carried out according to protocols approved by the Animal Care and Use Committee of the University of Pennsylvania. All experiments were performed on C57BL/6J mice, purchased from Jackson Laboratory. The mice were verified to be free of *Enterobacteriaceae* by plating stool on MacConkey agar. Female mice were 9 weeks old, male mice were 7 weeks old. For studies with antibiotic pre-treatment (Figures 1C,D and S2), mice were administered streptomycin (5 g/liter) and glucose (5 g/liter) in the drinking water for 72 hours; twenty-four hours after antibiotics were stopped, the mice were gavaged with bacterial suspensions. For experiments without antibiotics (Figures 1 A, B, E, F and 2), mice were allowed to acclimate for 96 hours prior to inoculation. To induce acute intestinal inflammation, mice were administered 4% Dextran Sodium Sulfate (DSS) (M.W.: 40,000 - 50,000; Affymetrix) in the drinking water as indicated in the timelines of Figure 1. During the DSS treatment, mice were monitored every day for the following: weight, general signs of distress, consistency of stool, and presence of blood in the stool. Occult bleeding was detected using the Hemoccult Screening Test and Hemoccult Sensa (Beckam Coulter, Brea, CA). Experiments were carried out according to the timelines reported in Figure 1, or until the disease activity index (DAI) (26) reached a value requiring euthanasia. Criteria for euthanasia were any one of the following: weight loss of 20%, DAI of 5, a score of 3 in one single category, or visible signs of distress (mice hunched over, lethargic, not eating or drinking)

To prepare bacterial suspensions for gavage, strains were grown overnight in LB. The culture optical densities were measured to normalize the inocula, and the appropriate culture volumes were centrifuged, resuspended in 5 ml ice-cold sterile phosphate-buffered saline (PBS), centrifuged again, and resuspended sterile PBS. For co-colonization experiments, equal volumes of the two competing strains were mixed, and 100 µl of this mixture was administered to mice by oral gavage. Mice inoculated with mono-cultures also were gavaged with 100 µl of bacterial suspensions. Each mouse received ~$10^{10}$ bacteria, except in experiments shown in Figures 1A and 1B, where each mouse received ~$10^{9}$ bacteria, as in (10).

To determine stool bacterial counts, fresh stool, or colon and cecum content post-euthanasia, were collected from each individual mouse, weighed and resuspended in sterile PBS at 250 mg of specimen per ml. The homogenized stool was serially diluted, and appropriate dilutions were spread on LB agar containing tetracycline to induce expression of the fluorescent proteins GFP and mCherry (Lasaro et al, 2014). After incubation at 37 °C, plates were imaged with a fluorescence illuminator, and green- and red-fluorescent colonies were counted.

Representative samples were collected for histopathological evaluation: colon was collected from mice, butterflied, trimmed, rolled and placed in histology cassettes. Tissue samples were fixed in 10% neutral buffered formalin (StatLab, MCKinney, TX) for 24 hours, washed twice in PBS, and stored in 70% ethanol until embedded in paraffin. Sections were stained with Hematoxylin and Eosin, blinded, and evaluated by a veterinary pathologist.

*In Vitro Studies*

To assess anaerobic respiration, strains were cultured overnight in M9 with 0.2% glucose, aerobically. The cultures were washed twice in sterile PBS and were diluted 1/1000 in M9 with glycerol and with or without the addition of 40 mM $KNO_3$ as indicated. The cultures were incubated in sealed screw-cap vials at 37 $^{o}$C for 24 hours, after which the culture optical densities at 600 nm were measured.

**Table S1. List of strains used in this study**

| Strain | Relevant Genotype | Source or reference |
| --- | --- | --- |
| JW0764 | BW25113 Δ*moaA*::[FRT-*kan*-FRT] | (27) |
| MP1 | *E. coli* Commensal isolated from mouse stool | (14) |
| MP7 | MP1 $att_{\lambda}$:: [*cat tetR* Φ(*tetA-mcherry*)] | (14) |
| MP13 | MP1 $att_{\lambda}$:: [*cat tetR* Φ(*tetA-gfp*)] | (14) |
| MP240 | MP1 Δ*moaA*::[FRT-*kan*-FRT] | This study |
| MP241 | MP13 Δ*moaA*::[FRT-*kan*-FRT] | This study |
| MP302 | MP13 Δ*moaA*::FRT | This study |
| MP321 | MP13 Δ*moaA*::FRT $att_{Tn7}$::*moaA* | This study |

**Table S2. List of primers used in this study**

| Primer | Sequence | Target Construct |
| --- | --- | --- |
| *moaA*-KEIO-F | CTCTGCACCTGGGTCAACTG | Δ*moaA*-MP1 |
| *moaA*-lower | GTGACCGGAGGTATCGTCTTC | Δ*moaA*-MP1 |
| *moaA*-XmaI-U1 | CCTGACCCCGGGCGATACCCGTTAAACGAGAACC[a] | *moaA*-MP1 complement |
| *moaA*-EcoRI-L1 | CCTGACGAATTCTTGACGTTTTAGCCACCAATG[a] | *moaA*-MP1 complement |

[a] Underlined sequence corresponds to the added restriction site.

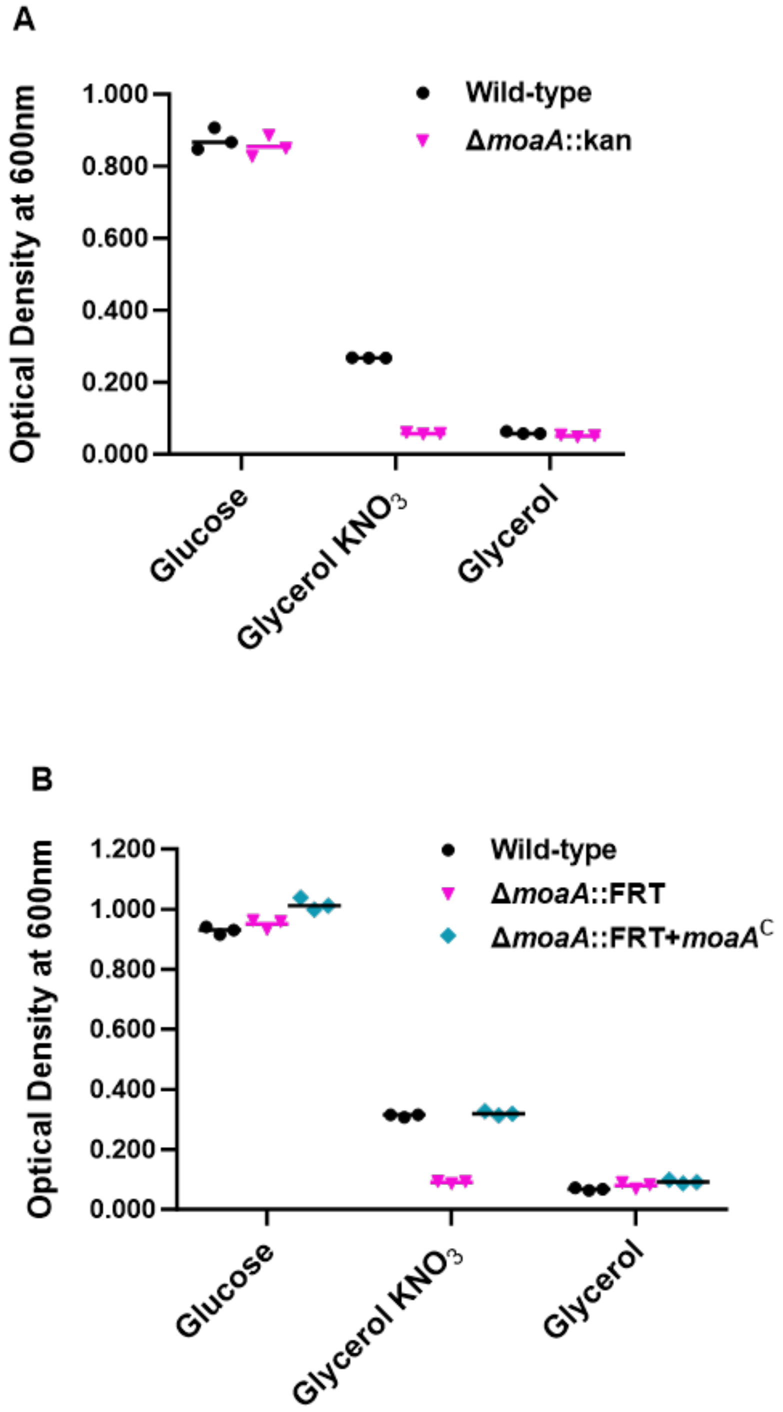


**Figure S1.** Cultures were grown for 24 hours anaerobically in minimal media with the indicated carbon sources and, where indicated, with potassium nitrate. The notation *moaA*$^{C}$ indicates the *moaA* gene was inserted at the *att*$_{Tn7}$ site.

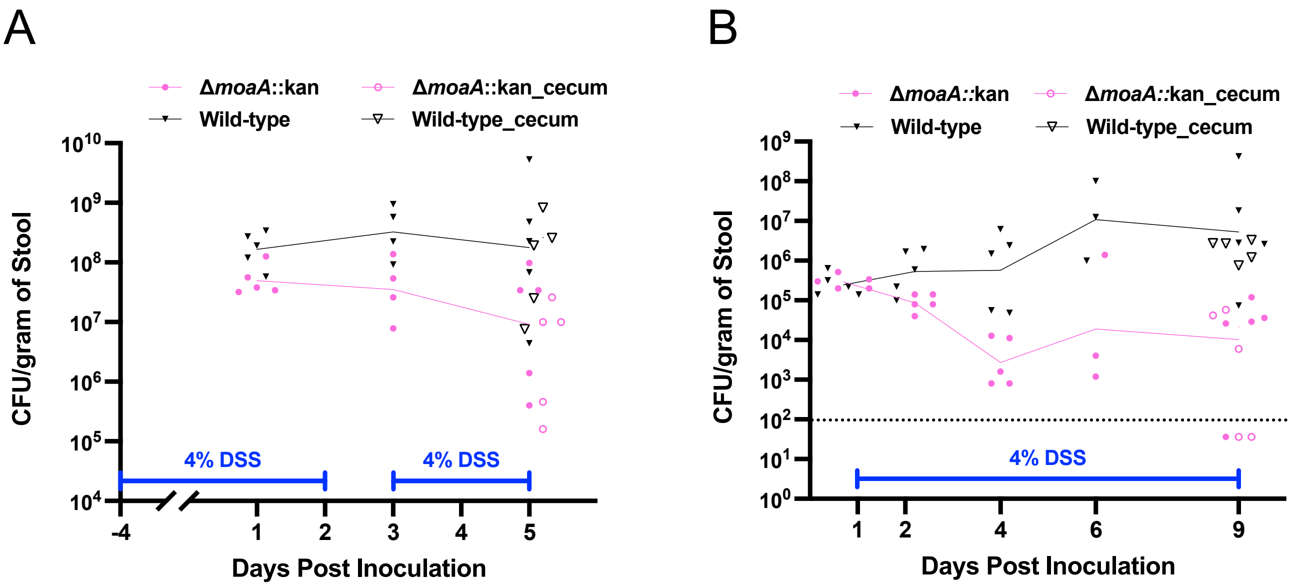


**Figure S2.** Stool CFU for the experiments shown in Figs. 1A,B. The dotted line in (B) denotes the limit of detection.

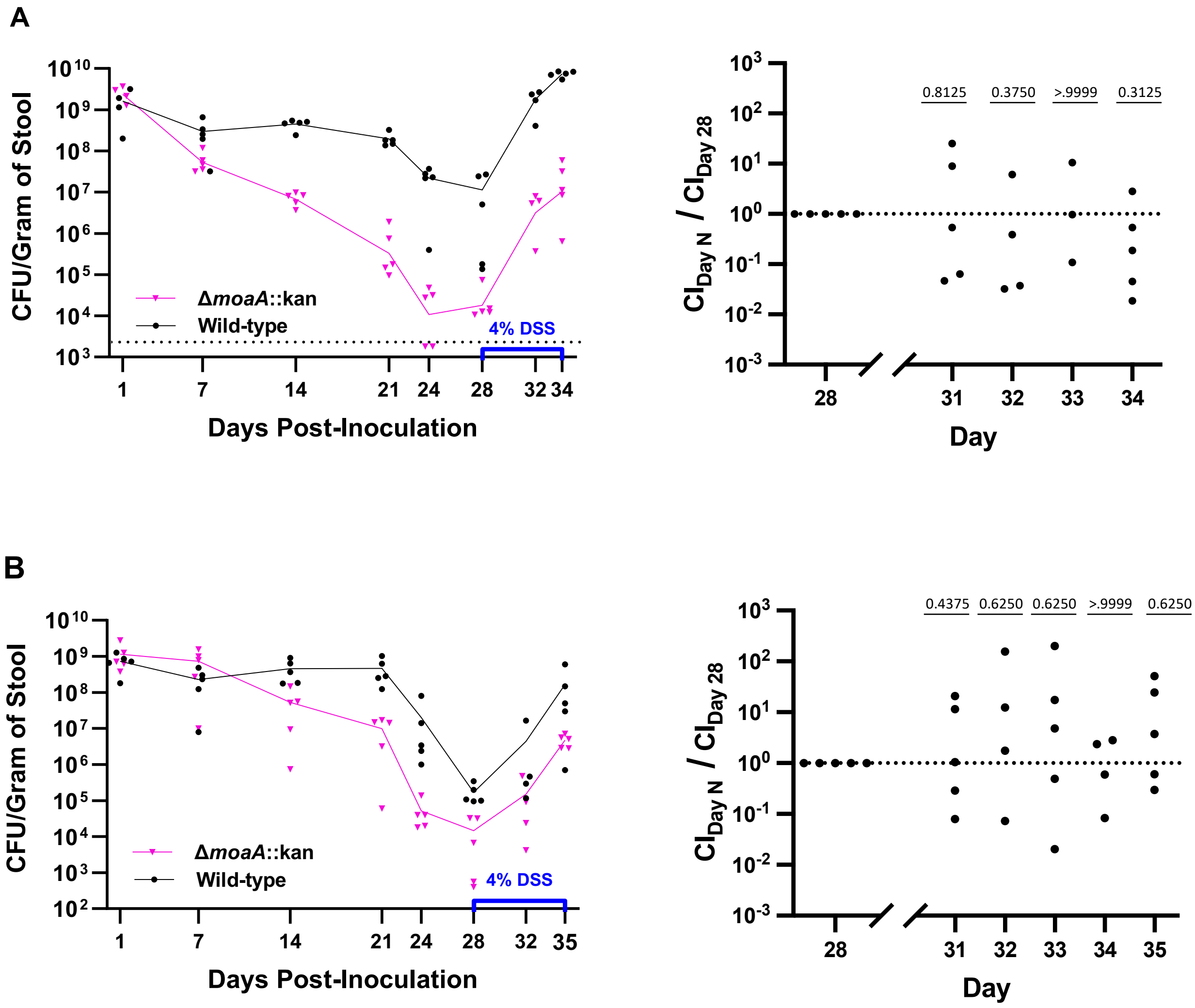


**Figure S3.** Panels (A) and (B) are two repeats of the mouse competition experiments shown in (Fig. 1 C,D); all three experiments were performed on different dates. For these two repeats, on day 21, mice were gavaged with a slurry of feces from untreated mice to bring down the CFU (see main text), and DSS was started on day 28. In the left panels of (A) and (B), points denote the CFUs from individual mice, and lines connect the geometric means. The dotted line in the left panel of (A) denotes the limit of detection. The right panels of (A) and (B) show the relative fold changes in abundance following DSS treatment for the data in the corresponding left panels. The relative fold change on day N of DSS treatment is defined to be the ratio of the competitive index, CI, on day N to the CI just prior to DSS treatment on day 28: $\mathrm{CI}_{\mathrm{Day\ N}}/\mathrm{CI}_{\mathrm{Day\ 28}}$ . Note that this ratio satisfies $\mathrm{CI}_{\mathrm{Day\ N}}/\mathrm{CI}_{\mathrm{Day\ 28}} = \frac{\left(\Delta moaA\ \mathrm{CFU}/\mathrm{WT\ CFU}\right)_{\mathrm{Day\ N}}}{\left(\Delta moaA\ \mathrm{CFU}/\mathrm{WT\ CFU}\right)_{\mathrm{Day\ 28}}}$ . The numbers above the data points in the right panels denote p-values from a Wilcoxon matched-pairs signed-rank test comparing wild type and Δ*moaA*. All DSS-treated mice showed evidence of weight loss and disease activity indices of at least 3 by the end of the experiments. The experiment in panel A was stopped after 6 days on DSS because the disease activity indices reached levels that required euthanasia. For each of the two experiments shown here, the five mice were co-housed in the same cage.

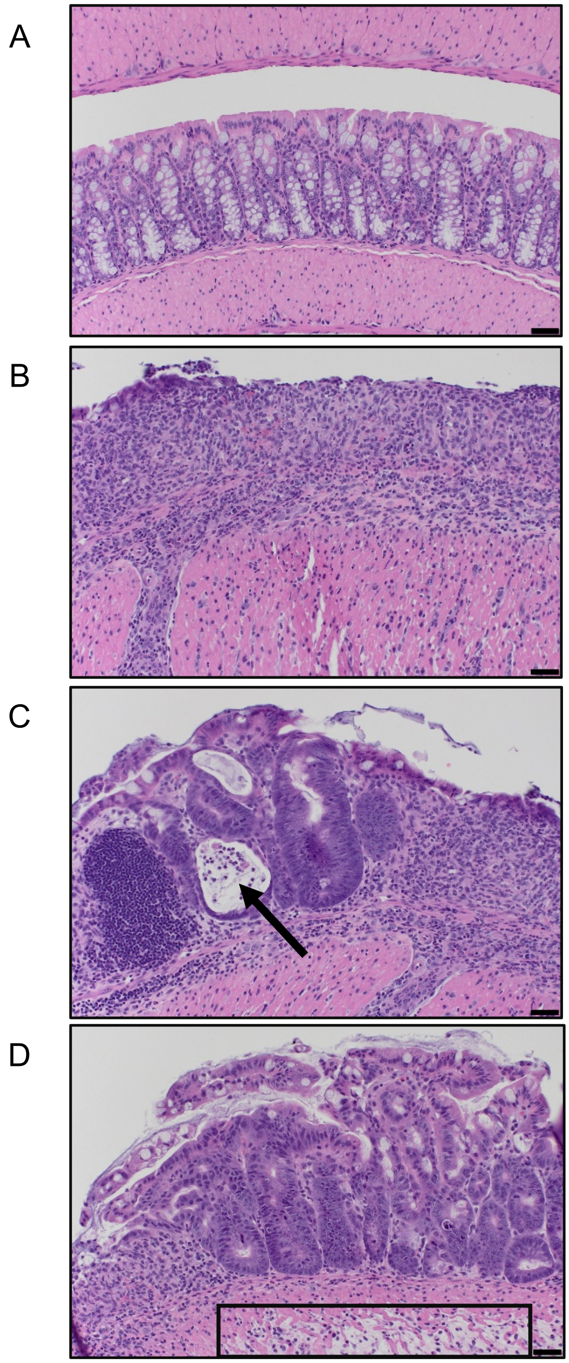


**Figure S4.** Representative images of H&E-stained colonic sections of (A) an untreated mouse and (B, C, D) a DSS-treated mouse (from Figure 1). The arrow in (C) points to a crypt abscess, and the box in (D) indicates edema.  Scale bar, 20 μm.
